# An *in vivo* avian model of human melanoma to perform rapid and robust preclinical studies

**DOI:** 10.1101/2022.10.12.511927

**Authors:** Loraine Jarrosson, Stéphane Dalle, Clélia Costechareyre, Yaqi Tang, Maxime Grimont, Maud Plaschka, Marjorie Lacourrège, Romain Teinturier, Myrtille Le Bouar, Delphine Maucort-Boulch, Anaïs Eberhardt, Valérie Castellani, Julie Caramel, Céline Delloye-Bourgeois

## Abstract

Metastatic melanoma patients carrying a BRAF^V600^ mutation can be treated with BRAF inhibitors (BRAFi), in combination with MEK inhibitors (MEKi), but innate and acquired resistance invariably occurs. Resistance can involve transcriptional- and epigenetic-based phenotypic adaptations, as yet unpredictable. Predicting patient response to targeted therapies is crucial to guide clinical decision. We describe here the development of a highly efficient patient-derived xenograft model adapted to patient melanoma biopsies, using the avian embryo as a host (AVI-PDX^™^). In this *in vivo* paradigm, we depict a fast and reproducible tumor engraftment of patient samples within the embryonic skin, preserving key molecular and phenotypic features. We show that sensitivity and resistance to BRAFi/MEKi targeted therapies can be reliably modeled in these AVI-PDX^™^, as well as synergies with other drugs, such as HDACi. We further provide proof-of-concept that the AVI-PDX^™^ models the diversity of responses of melanoma patients to BRAFi/MEKi, within days, hence positioning it as a valuable tool for the design of personalized medicine assays and for the evaluation of novel combination strategies.

## Introduction

Cutaneous malignant melanoma is an aggressive form of skin cancer arising from melanocytes. Despite recent advances in targeted therapies and immunotherapies for the treatment of metastatic melanoma, nearly 60% of patients still develop resistance, necessitating the development of new therapeutic strategies in mono- or combination-therapy (Luke *et al*, 2017; Herrscher & Robert, 2020; Dummer *et al*, 2020; Curti & Faries, 2021). Metastatic melanoma (MM) patients carrying a BRAF^V600^ mutation (50% of cases) can be treated with BRAF inhibitors (BRAFi), in combination with MEK inhibitors (MEKi), but innate (40%) or acquired resistance invariably occurs (Larkin *et al*, 2014; Trunzer *et al*, 2013).

Increasing evidence suggests that resistance to BRAFi/MEKi is not only mediated by genomic alterations, but also involves phenotypic adaptations through transcriptional and epigenetic processes (Hugo *et al*, 2016; Rambow *et al*, 2018; Marine *et al*, 2020). Cellular plasticity achieved through epithelial-to-mesenchymal (EMT)-like processes contributes to intra-tumor heterogeneity (ITH) in melanoma and fosters the ability of cancer cells to adapt to treatment (Rambow *et al*, 2019; Tang *et al*, 2020). Such reversible phenotypic transitions between a proliferative/differentiated and invasive/stem-like state (Hoek *et al*, 2008), are reminiscent of the features acquired upon delamination of the embryonic neural crest from which melanocytes originate (Mort *et al*, 2015). Indeed, melanoblasts originate from neural crest cells (NCCs) that transit through a SOX10 (Sry-related HMG-box-10)-positive melanoblast/glial bipotent progenitor state. Specified melanoblasts acquire MITF (microphthalmia-associated transcription factor) expression, mostly migrating dorsolaterally from the dorsal neural tube to reach the skin, where they differentiate into melanocytes that produce the melanin pigment. MITF, the master regulator of melanocyte differentiation, is also a major regulator of melanoma phenotype switching (Goding & Arnheiter, 2019). Loss of MITF induces a reprogramming towards an invasive and stem-like phenotype in melanoma cells (Goding & Arnheiter, 2019; Rambow *et al*, 2019). Reprogramming towards a neural crest stem cell-like (NCSC) phenotype, via the re-expression of NCSC markers (such as NGFR), was proposed as an adaptive response to targeted therapy, accounting for therapy resilience (Diener & Sommer, 2021). The EMT-inducing transcription factor ZEB1 also promotes the transition towards a stem-like and invasive MITF^low^ state, resistant to BRAFi/MEKi therapy (Caramel *et al*, 2013; Richard *et al*, 2016). While biomarkers of response to targeted therapies have been proposed, robust tools able to predict patient response in a time-frame compatible with the clinical decision, constitute unmet medical needs.

Various *in vivo* melanoma models (mouse, fish) have been developed over the years (Patton *et al*, 2021), each presenting pros and cons with respect to their capacity to reproduce the human disease, the tumor heterogeneity, the tumor microenvironment (TME), and to provide relevant results for chemical screens. Starting from fresh human samples, despite their preclinical relevance, patient-derived xenograft models (PDXs) in the mouse, are hampered by their high-cost and long timeframe for evaluating treatment efficacy, limiting their application in personalized medicine programs. Organoids may prove useful for high-throughput chemical screens, but they lack crucial components of the TME and have so far not been extensively validated for melanoma (Ronteix *et al*, 2022). Testing drug efficacy on *ex vivo* tumor fragments was recently shown to hold a robust predictive capacity, including for immunotherapy (Voabil *et al*, 2021). However, it requires a significant tumor size, incompatible with large drug screenings and with statistical evaluation of drug anti-tumor efficacy. The development of a miniaturized PDX model, requiring low amounts of tumor samples and displaying a short-timeframe (a few days) of development, would thus be a valuable preclinical tool for assessing treatment efficacy in melanoma. We recently described the development of an animal model combining these key advantages for neuroblastoma and triple negative breast cancers (TNBCs), using the avian embryo as a recipient organism for patient graft samples (Delloye-Bourgeois *et al*, 2017; Jarrosson *et al*, 2021).

Herein, starting from melanoma cell lines (BRAFi/MEKi sensitive/resistant) and then from human melanoma samples, we provide the proof-of-concept for the development of an innovative, highly efficient and reproducible melanoma PDX model using the avian embryo as a host. We show that this miniaturized paradigm which reproduces melanoma cell phenotypes in their microenvironment, allows to assess drug combinations, and to mimic MM patient clinical responses to targeted therapies. As such, the avian embryo PDX model is ideally suited for testing patient tumor responses in a timeframe compatible with therapeutic decision-making by clinicians.

## Results

### Micrografting human melanoma cell lines in the neural crest of the avian embryo

With the aim of designing a melanoma model mimicking the localization of primary lesions in the skin and their eventual secondary dissemination, we sought to target the original lineage of melanocytes, the neural crest cells. Indeed, during development, NCCs emerge from the dorsal roof of the neural tube along the rostro-caudal axis. A subset of these cells reach the so-called migration staging area (MSA), in which they become melanoblasts, before engaging into a stereotyped dorsolateral trajectory to ultimately populate the skin and hair follicles (Mort *et al*, 2015). We hypothesized that placing melanoma cells among the pre-migratory melanoblast cell population, within the MSA, could offer a supportive microenvironment to foster their migration and establishment in the skin. Thus, we engrafted human melanoma cells (BRAF^V600E^-mutated A375P cell line) engineered to stably express GFP (A375P::GFP) next to the dorsal roof, at the MSA level, in HH13 (according to Hamburger and Hamilton staging method) chick embryos, in a trunk region lying between somite 16 and somite 24 (**Fig. 1A**). Forty-eight hours after engraftment, embryos were collected at the HH25 stage and the position of cells was analyzed by 3D imaging using lightsheet confocal microscopy and by immunofluorescent staining on transverse sections. 3D analysis of engrafted embryos revealed that, in all embryos, A375P::GFP cells were no longer present at the graft site but had diffused under the skin and in deeper tissues (**Fig. 1B**). Previous work reported that when engrafted at earlier stages of chick development, melanoma cells spontaneously differentiate and lose their tumorigenic properties (Kulesa *et al*, 2006). We thus assessed whether in our paradigm grafted cells maintained their proliferative state. A strong Ki67 immunofluorescent labeling was observed in A375P::GFP cells within the tumor mass highlighting high proliferative activity (**Fig. 1C**). We observed that grafted cells did not migrate randomly but were localized within typical streams of NCCs dorsolateral and ventrolateral migration paths, subsequently reaching the developing skin or under the epidermis, but not in developing sensory and sympathetic ganglia (**Fig. 1BC**). Hence, the neural crest origin of these melanoma cells may have impacted their ability to integrate microenvironmental signals, which are known to influence NCC migratory and differentiation properties. We then wondered whether exposure of melanoma cells to this particular embryonic microenvironment could have modified the expression of key actors of the SOX10-MITF axis, well described to be involved in melanomagenesis (Capparelli *et al*, 2022; Shakhova *et al*, 2015; Cheli *et al*, 2011). A375P cells are known to display a NCSC-like expression pattern (MITF^low^, SOX10^+^) as recently analyzed at the single cell level (Wouters *et al*, 2020). Consistently, immunofluorescence analyses of A375P cells engrafted in the avian embryo revealed a heterogeneous expression of MITF with typically low or negative cells, and a few positive cells, while SOX10 expression (also detected in chick endogenous neural crest cells) was homogeneous, suggesting preservation of molecular features involved in their tumorigenic properties (**Fig. 1D**). Thus, implantation of human melanoma cells in a selected embryonic stage and territory drives tumor growth in relevant tissues and maintenance of their tumorigenic potential.

**Figure 1:**
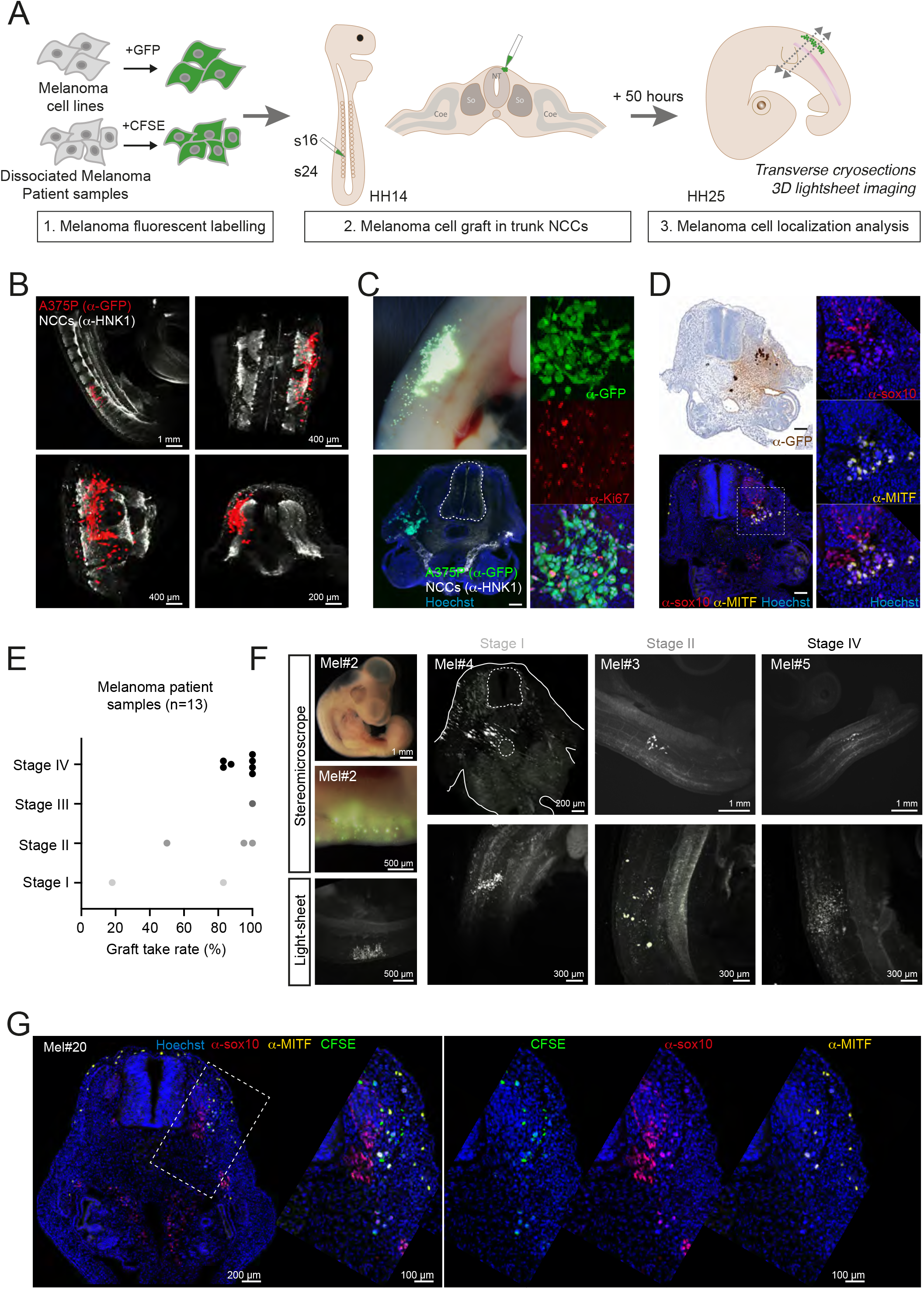
Set up and characterization of melanoma cell lines and patient samples engrafted in the avian embryo. **A.** Schematic diagram of the engrafting procedure of melanoma cell lines or melanoma biopsies in the chick embryo. **B.** 3D views (light-sheet imaging) of HH25 chick embryos engrafted with the A375P::GFP stable melanoma cell line, immunolabeled with an anti-GFP antibody (in red, A375P:GFP cells) and an anti-HNK1 antibody (in white, migrating and early post-migrating NCCs). **C,D.** Immunolabeling of HH25 chick embryo sections 50 hours post-engraftment of A375P::GFP cells, using an anti-GFP antibody (in green in **C**, in brown in **D**, A375P:GFP cells), an anti-HNK1 antibody (in white, migrating and early post-migrating NCCs), an anti-Ki67 antibody (in red in **C**, cycling cells), an anti-SOX10 antibody (in red in **D**) and an anti-MITF antibody (in yellow in **D**). Nuclei were stained with Hoechst (in blue). In **C**, the upper left photo shows a grafted embryo prior to cryosection. In **D**, right panels are enlargements of the lower left panel. Scale bar: 200 μm. **E.** Tumor take rate of 13 melanoma patient samples engrafted in a series of chick embryos, ranked according to their assigned Stage (I to IV). **F.**3D views (light-sheet imaging) of HH25 chick embryos engrafted with different melanoma patient samples, showing different patterns of tumor cell localization 50 hours post-engraftment. **G.** Immunofluorescent labeling of HH25 chick embryo sections 50 hours post-engraftment of OF-MEL-020 patient sample, labeled with CFSE (in green) prior to the graft. An anti-SOX10 antibody (in red) and an anti-MITF antibody (in yellow) were used. Nuclei were stained with Hoechst (in blue). Right panels are enlargements of the dotted area in the left panel.

### Generation of patient-derived xenograft melanoma models using the avian embryo as a host (AVI-PDX™)

To extend the preclinical applicability of our *in ovo* micrografting model, we examined the behavior of melanoma cells isolated from patient samples following the same procedure as for cell lines. Thirteen melanoma biopsies from primary or metastatic melanoma (Supplementary Table 1), were enzymatically dissociated and labelled with Carboxyfluorescein succinimidyl ester (CFSE) to trace their behavior after engraftment in a series of avian embryos (Fig. 1E). Remarkably, successful tumor graft was observed for each patient, even for lower stages tumors, highlighting a highly robust model to sustain tumor growth of human melanoma samples (Fig 1E). The tumor graft take rate 48 hours following transplantation was above 50% for all patient samples, except for the OF-MEL-001 patient showing a very low cellularity (18% of tumor take). Consistently with what was previously observed for human TNBC samples in the avian graft paradigm (Jarrosson *et al*, 2021), we did not notice any significant correlation between tumor stage or proliferative index (nb of mitosis/mm2) and tumor take rate (**Supp. Table 1**).

Analysis of embryos by 3D light-sheet confocal microscopy revealed different distribution patterns of patient cells within the embryonic tissues ranging from scattered cells with few cell-cell contacts to dense, cohesive tumor foci under the skin (**Fig. 1F**). Notably, such patterns were reproduced in all embryos engrafted with the same patient sample. As for the A375P cell line, we performed SOX10 and MITF labelling on cryosections of embryos engrafted with patient samples. We confirmed that a fraction of CFSE-labelled tumor cells also expressed varying levels of MITF and/or SOX10 along their migration path, indicating that melanoma intra-tumor heterogeneity (ITH) is preserved after implantation in the embryonic environment (**Fig. 1G**). Altogether, the AVI-PDX^™^ constitutes a highly efficient and reproducible approach to create PDX models of human melanoma.

### Proliferative and invasive melanoma phenotypes are preserved upon grafting in the avian model

Since ITH is a key feature of melanoma, participating in resistance to therapies, particularly to BRAF/MEK inhibitors (BRAFi/MEKi) (Marine *et al*, 2020), we assessed whether the dynamic phenotypic switches underlying melanoma cell proliferative/ invasive states could be modeled in the AVI-PDX^™^. We initially analyzed the expression profiles of MITF and Ki67 in A375P-xenografts, in which dense tumor masses were surrounded by streams of migrating cells (**Fig. 2A,B**). Interestingly, the fraction of A375P cells showing no or low MITF expression was significantly higher in cells disconnected from dense tumor masses, suggesting a dynamic regulation of MITF expression depending on the proliferative (tumor masses) versus invasive (migrating cells) states of melanoma cells (**Fig. 2A,B**). Consistently, the fraction of Ki67-negative cells followed the same tendency, with melanoma cells with a typical mesenchymal morphology being negative for Ki67 (**Fig. 2C,D**).

**Figure 2:**
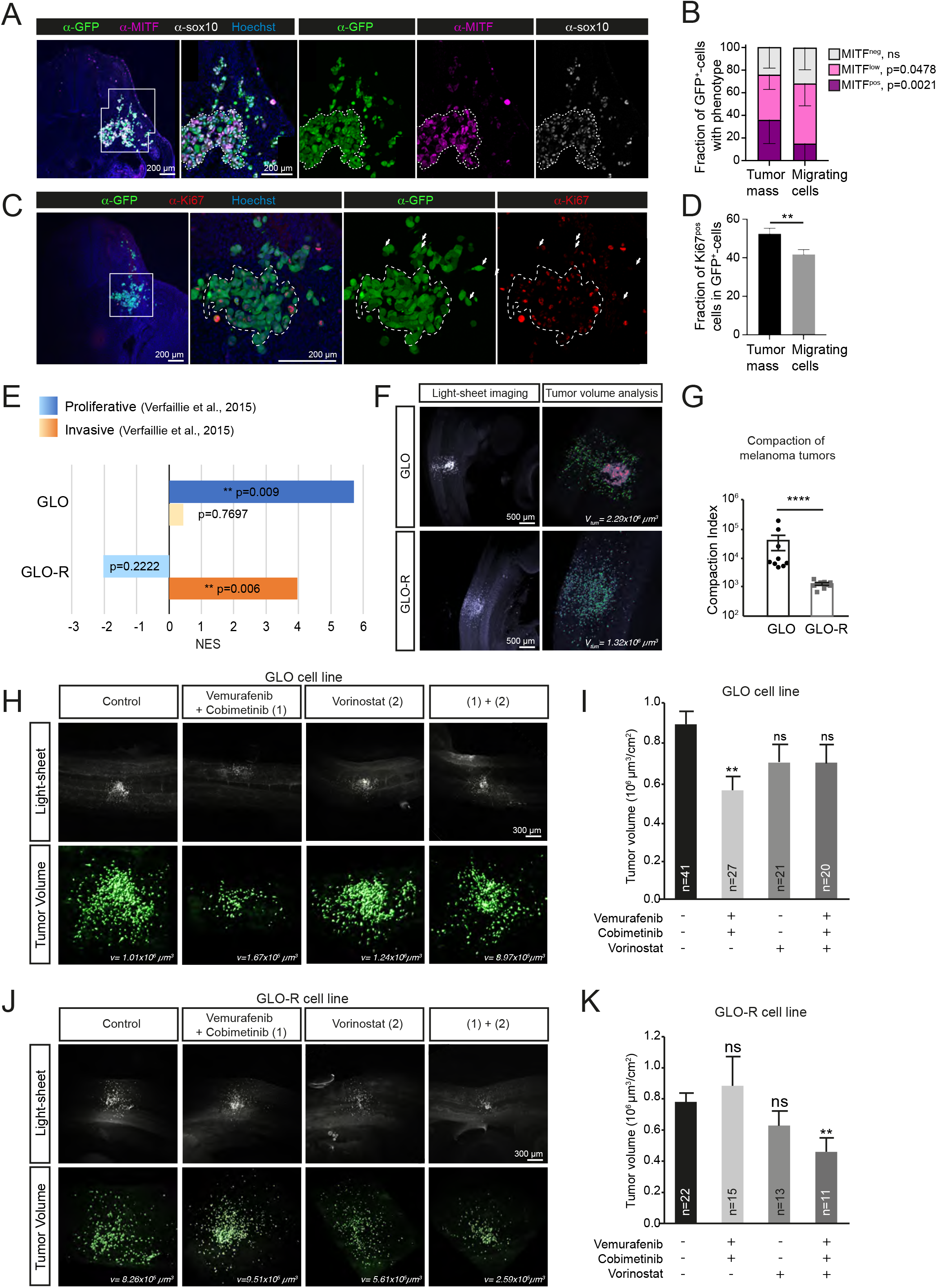
Melanoma cells retain their phenotypic traits and associated response to targeted therapies in the AVI-PDX™ model. **A,B.** Immunofluorescent labeling (**A**) of HH25 chick embryo sections 50 hours post-engraftment of A375P::GFP cells, using an anti-GFP antibody, an anti-MITF antibody (pink) and an anti-SOX10 (white) antibody. Nuclei were stained with Hoechst. Right panels are enlargements of the dotted area in left panel. In **B**, the mean fraction of MITF-negative (MITF^neg^),-low (MITF^low^) and -positive (MITF^pos^) A375P::GFP cells in the tumor mass and in migrating cells was quantified in n = 16 slices. Error bars show SEM. Mann-Whitney test comparing tumors *vs* migrating cells for each class of MITF expression was performed, p-values are indicated on the graph. **C,D.** Immunofluorescent labeling (**C**) of HH25 chick embryo sections 50 hours post-engraftment of A375P::GFP cells, using an anti-GFP antibody and an anti-Ki67 antibody. Nuclei were stained with Hoechst. Right panels are enlargements of the dotted area in the left panel. In **D**, the mean fraction of Ki67-positive (Ki67^pos^) A375P::GFP cells in the tumor mass and in migrating cells was quantified in n = 25 slices. Error bars show SEM. Student t-test, p = 0.0070. **E.** RNASeq analysis of GLO and GLO-R cells; gene signatures published in (Verfaillie *et al*, 2015), associated with either a proliferative or an invasive melanoma phenotype were scored in both cell lines. P-values are indicated in the graphical representation. **F,G.**3D views (light-sheet imaging, **F**) of HH25 chick embryos engrafted with GLO or GLO-R cells labeled with CFSE (in green) prior to the graft. The total volume occupied by tumor cells and the number of segmented CFSE-positive objects was quantified in **G** to calculate a mean compaction index (see details in the Materials and Methods section) of tumors for each cell line. Mann-Whitney test, p < 0.0001. **H-K**. 3D views (**H, J**) and quantification of tumor volumes (**I, K**) of HH25 chick embryos engrafted with GLO (**H, I**) or GLO-R (**J, K**) cells and treated with a combination of Vemurafenib and Cobimetinib, or Vorinostat alone, or a combination of the three molecules. The number of embryos analyzed for each experimental condition are indicated on the graphs. Student t-tests, **: p < 0.01, ns: not significant.

Next, based on recently established patient-derived short-term cultures we investigated the possibility of mimicking melanoma cell phenotypes and associated clinical responses to BRAFi/MEKi. We used the GLO and GLO-R pair of BRAFi/MEKi sensitive/resistant primary cell lines, established from the same BRAF^V600^-mutated MM patient, before (GLO) (Richard *et al*, 2016) or after (GLO-R) the emergence of resistance to BRAFi/MEKi. These clinical profiles were reproduced *in vitro* by measuring cell survival upon treatment with increasing doses of vemurafenib or cobimetinib (**Supp. Fig. 1A,B**). Western blots of these two patient-derived melanoma cultures revealed that both MITF and SOX10 expression were decreased in BRAFi/MEKi-resistant GLO-R cells (**Supp. Fig. 1C**). In accordance with previously published data (Richard *et al*, 2016), the levels of the EMT-TF, ZEB1, were higher in GLO-R compared to the BRAFi/MEKi-sensitive GLO line, while ZEB2 expression levels decreased. To more precisely document the changes occurring in patient cells upon acquisition of BRAFi/MEKi-resistance, we performed a comparative bulk RNA-Seq analysis of GLO and GLO-R cell lines (**Fig. 2E**). We assessed the expression of gene signatures typically involved in melanoma cell plasticity. Interestingly, while GLO cells displayed an enrichment of genes involved in cell proliferation, GLO-R cells were characterized by an invasive signature, evoking a switch towards an aggressive, motile, cell behavior upon acquisition of resistance (**Fig. 2E**).

We then assessed the behavior of GLO and GLO-R primary cell lines when engrafted in the avian embryo. We performed 3D reconstruction of the tumors using confocal light-sheet microscopy (**Fig. 2F**). Interestingly, the distribution pattern of cells in the embryonic tissues was different between the two cell lines. While GLO cells were detected as cohesive tumor masses immediately surrounded by isolated migrating cells, individual GLO-R cells were scattered under the skin and in deeper tissues, rarely forming clusters. Quantification of tumor cell dispersion achieved by measuring the compaction index (as detailed in the methods section) confirmed a significant difference between GLO and GLO-R tumor features (**Fig. 2G**). Thus, the divergent proliferative and invasive states of GLO and GLO-R cells were reflected in their distinct tumor patterns within embryonic tissues.

Moreover, immunolabelling of transverse sections of avian embryos showed that engrafted GLO cells maintained a high level of MITF and SOX10 expression compared to GLO-R cells, in which MITF expression was negligible (**Supp. Fig. 1D**). These data confirm that key molecular features of the SOX10-MITF axis are preserved 48 hours post-engraftment in chick embryos and suggest that differences in GLO/GLO-R invasive capacities mirrored their respective expression levels of MITF.

Thus, by recapitulating the embryonic microenvironment and preserving key intrinsic molecular features, the AVI-PDX^™^ enables melanoma cells to translate their different SOX10-MITF levels into distinct migratory/invasive behaviors.

### The avian melanoma model robustly predicts drug efficacy in melanoma patients

We then investigated whether our AVI-PDX^™^ could reproduce the heterogeneity of patient tumor responses to BRAFi/MEKi therapies, to offer a novel solution for preclinical studies in melanoma. We further evaluated the efficacy of these therapies in combination with epigenetic drugs, namely HDAC (histone deacetylases) inhibitors such as Vorinostat, which emerged as therapeutic options to overcome resistance, showing promising results in preclinical and clinical studies (Huijberts *et al*, 2020).

We initially determined the optimal dose of each therapeutic compound-i.e., Vemurafenib, Cobimetinib, Vorinostat-in chick embryos. Having access to the chorioallantoic membrane that irrigates the developing embryo, we performed intravenous injections of increasing doses of each drug separately, in series of HH20 chick embryos (approximately 72 hours post-gestation) (**Supp. Fig1E-G**). Twenty-four hours post-injection (HH25 embryos), we quantified the survival rate of injected embryos. We also assessed their development by examining key morphological checkpoints, as described in the methods section and in previous work (Jarrosson *et al*, 2021), and measured global growth by quantifying the body surface area (BSA) of each viable embryo. Survival rates below 75% or significant differences in BSA compared to the control group (treated with the excipient of each drug) were indicative of dose toxicity. According to these criteria, the *in ovo* maximum tolerated dose (MTD) of Cobimetinib, Vemurafenib and Vorinostat were defined at 0.0036 mg/kg, 13.1 mg/kg and 0.744 mg/kg, respectively. We then assessed whether administration of Cobimetinib/Vemurafenib combitherapy, Vorinostat or combination of the three drugs could impair on GLO and GLO-R tumor growth after their engraftment *in ovo* (**Fig. 2H-K**). Tumor volumes were measured by 3D light-sheet confocal imaging of whole chick embryos after a 24 hour-treatment. Consistent with our *in vitro* data, co-injection of Vemurafenib and Cobimetinib triggered a significant reduction of GLO-tumor volumes compared to excipient-treated embryos (**Fig. 2H,I**) while GLO-R-tumor volumes were not affected (**Fig. 2J,K**). Hence, the sensitivity/resistance features of BRAF^V600^-mutated melanoma cells to BRAFi/MEKi were maintained in the avian model. Vorinostat alone did not impact GLO-tumor volumes and only triggered a slight decrease in size of BRAFi/MEKi resistant GLO-R tumors. Conversely, co-injection of Vorinostat with Vemurafenib/Cobimetinib triggered a significant reduction of GLO-R tumor volumes, which is in favor of recent data suggesting that BRAFi/MEKi resistance could, at least partially, be overcome by epigenetic drugs (Wang et al., 2018) (**Fig. 2J,K**). Notably, when Vorinostat was combined with BRAFi/MEKi, the anti-tumor effect of BRAFi/MEKi on GLO-tumors was abrogated (**Fig. 2H,I**). This observation corroborates previous studies suggesting that HDACi could antagonize BRAFi/MEKi activity in BRAFi/MEKi-sensitive melanoma cells (Wang *et al*, 2018). These findings suggest that our melanoma AVI-PDX^™^ model may be promising for preclinical studies on mono- and combination therapies.

### The AVI-PDX™ allows relevant preclinical assessment of targeted therapies and is predictive of patient clinical response

For further validation, we next evaluated the effect of Vemurafenib/Cobimetinib treatment on patient samples with distinct mutational profiles after their implantation in a series of chick embryos (**Fig. 3A,B**). Of note, a reduction of patient-derived tumors was observed upon Vemurafenib/Cobimetinib treatment in OF-MEL-027 and OF-MEL-020 samples, both harboring a BRAF^V600E^ mutation. Conversely, the NRAS^Q61L^-mutated OF-MEL-028 sample, having a wild type BRAF status, did not show any significant response to Vemurafenib/Cobimetinib treatment *in ovo*, in accordance with its mutational status.

**Figure 3:**
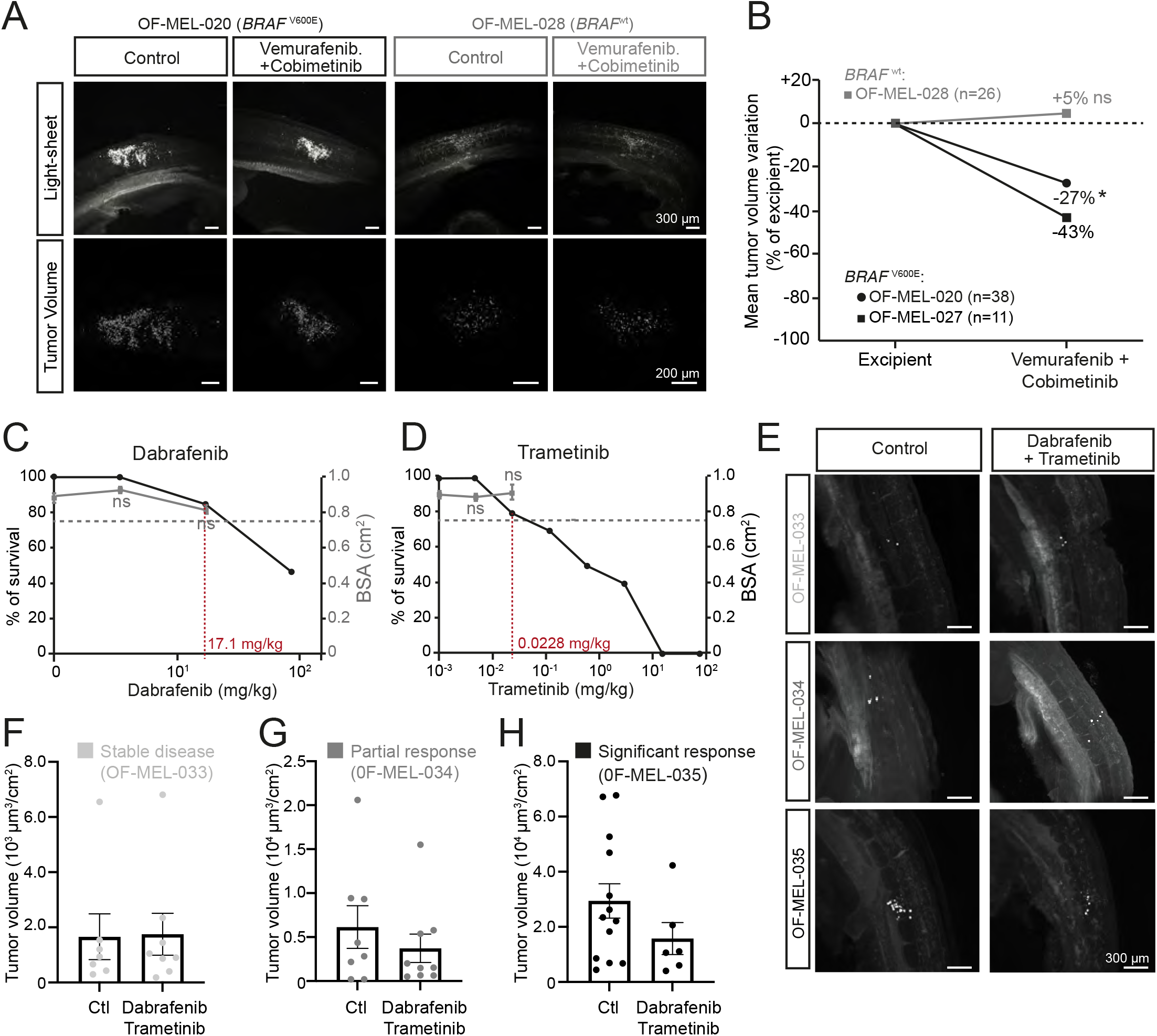
The AVI-PDX™ paradigm efficiently models patient clinical response to targeted therapies. **A,B.** 3D views (**A**) and quantification of variations in the mean tumor volume (**B**) of HH25 chick embryos engrafted with BRAF^wt^ (OF-MEL-028) or BRAF^V600E^ (OF-MEL-020, OF-MEL-027) patient samples and treated with excipient or a combination of Vemurafenib and Cobimetinib. The number of embryos analyzed for each patient sample is indicated on the graphs. Student t-tests, *: p < 0.05. **C,D**. Survival rate (left axis) and mean body surface area (BSA, right axis) of chick embryos injected with increasing doses of Dabrafenib (**C**) and Trametinib (**D**). Each dose was administered to a minimum of 10 embryos, using excipient (NaCl) as a control. The maximum tolerated dose (MTD) was defined as the higher dose of drug associated with a survival rate higher than 75% and a mean BSA similar (*i.e*., non-statistically different) from embryos treated with NaCl. MTDs are indicated in red on the X-axis. Error bars indicate SEM. ns, non-significant using Student’s t-test compared with excipient. **E-H.**3D views (**E**) and quantification of tumor volumes (**F-H**) of HH25 chick embryos engrafted with 3 different patient samples (OF-MEL-033 (**F**), OF-MEL-034 (**G**), OF-MEL-035 (**H**)) and treated with excipient or a combination of Dabrafenib and Trametinib. The clinical response of each patient after a 3 months treatment with Dabrafenib/Trametinib is indicated above the graphs.

We then studied whether the AVI-PDX^™^ model could be predictive of the clinical response of patients. In a prospective study, we compared the response of patients receiving BRAFi/MEKi within the first three months of treatment with that of their tumor replicas in the avian model. Three BRAF^V600E^-mutated patients, treated with a Dabrafenib/Trametinib combination, were clinically scored at three months which revealed three types of responses: stable disease (OF-MEL-033), partial response with local reduction of tumor foci (OF-MEL-034), or significant global response (OF-MEL-035). In parallel, replicas of these patient tumors were produced in avian embryos that were treated with the same molecules, Dabrafenib and Tramentinib, at their maximum tolerated dose (17.1 and 0.0228 mg/kg respectively) established as for Cobimetinib and Vemurafenib (**Fig. 3C,D**). The effect of Dabrafenib/Trametinib treatment on tumor volume was measured following the same method as described above (**Fig. 3E-H**). Remarkably, not only did the clinical stable disease evaluation match the stable volume of tumors in the avian replicas (OF-MEL-033, **Fig. 3F**), but the significant anti-tumor response to BRAFi/MEKi observed for patient OF-MEL-035 was also associated with a 44% decrease in tumor volume in avian replicas (**Fig. 3H**). Moreover, the partial response of patient OF-MEL-034 was translated into a discrete mean tumor volume reduction, with a strong heterogeneity of response between the tumor replicas (**Fig. 3G**). Thus, this analysis revealed a striking similarity between the clinical outcome of the patient and the short-term response of avian replicas.

## Discussion

Our study depicts an alternative *in vivo* model of melanoma and provides an overview of its power to perform relevant preclinical studies. When implanted at the level of the NCC migration staging area in HH14 chick embryos, we could show that melanoma cell lines, patient-derived short-term cultures but also fresh or frozen patient biopsies settled in the skin and under the epidermis, and formed tumor foci within 48 hours. Interestingly, grafted cells followed typical endogenous melanoblast migrating routes to reach the developing dermis and epidermis. There, tumor cells maintained their phenotypic heterogeneity and plasticity in terms of proliferative and invasive properties, and the expression of their corresponding key markers. Notably, tumor take was obtained for the 13 patient biopsies used in the study, and the tumor take rate was above 80% for 11 of them, irrespective of the stage, mitotic index, the metastatic/primary origin of the biopsy or the mutational status.

Melanoma patient-derived xenograft (PDX) models classically set up in mice are associated with major constraints among which the need for large amounts of tumor material incompatible with most melanoma biopsies, a very low graft take efficiency and a long-term establishment limiting statistical analyses and precluding studies designed for personalized medicine. Moreover, costs and ethical issues strongly limit mouse PDX applications. We show here that using the avian embryo as a host, melanoma patient biopsies can be implanted in a series of embryos without any culture step, leading to a miniaturized, fast, efficient and reproducible graft take in a relevant microenvironment that models melanoma cell heterogeneity and associated resistance/sensitivity profiles to targeted therapies. Of note, at developmental stages used herein, the use of chicken embryos is in perfect accordance with ethical guidelines according to the European directives 2010/63/EU.

Importantly, the results of our prospective analyses suggest that the effect of targeted therapies injected in melanoma AVI-PDX^™^ are predictive of immediate patient response to BRAFi/MEKi. The paradigm enabled us to characterize the response of a patient sample to targeted therapies within a timeframe compatible with therapeutic decision-making, suitable with a key criterion of personalized therapeutic care. Moreover, we could model synergistic/antagonistic behaviors of combination of targeted therapies depending on melanoma cell phenotypes, underlining the power of our in vivo paradigm to provide relevant information on candidate molecules that could be included in a complex therapeutic regimen. Overall, we provide proof-of-concept that the AVI-PDX^™^ model accurately reproduces melanoma patient response to BRAFi/MEKi, highly suited to assess the efficacy of drug combinations.

Immunotherapies targeting negative regulatory checkpoints in immune cells, are another active therapeutic option for patients with BRAF^V600^-mutant melanoma, though around 60% of patients still develop resistance to anti-PD-1 blocking antibodies therapy (Larkin *et al*, 2019). The most effective first-line treatment and the optimal sequencing of these agents is still matter of intense clinical research studies. Some studies suggest lower activity of immunotherapy after BRAFi/MEKi treatment (Simeone & Ascierto, 2017; Amini-Adle *et al*, 2018) and mechanisms of crossresistance between targeted and immunotherapies were recently characterized (Haas *et al*, 2021). But there are limited data on BRAFi/MEKi after immunotherapy failure (Xia *et al*, 2018; Rogala *et al*, 2022). Several clinical trials are ongoing to compare the efficacy of different sequences and regimens of BRAF/MEK therapy and immunotherapy and results are awaited.

The clinical decision about whether to first use targeted therapy or immunotherapy in patients with BRAF^V600^ MM is based mostly on the clinical characteristics but biomarkers are lacking. There is thus an urgent medical need to define which patients with BRAF^V600^ MM would benefit or not from a treatment with BRAF/MEK therapy and would better be directed to immunotherapy as a first-line treatment. In the future, our study should pave the way for the design of a test of personalized medicine, thus improving the therapeutic decision-making process for BRAF^V600^ melanoma patients.

## Material and Methods

### Anticancer drugs

Vemurafenib (PLX4032), Cobimetinib (GDC-0973), Vorinostat (SAHA), Dabrafenib (GSK2118436) and Trametinib (GSK1120212) were purchased from Selleckchem (stock solution at 10 mM). Those chemicals were diluted in DMSO - 0.5% Tween 80 used as an excipient, for *in vivo* experiments.

### Chick embryos

Embryonated eggs were obtained from a local supplier (Couvoir de Cerveloup, Vourey, France). Laying hen’s sanitary status was regularly checked by the supplier according to French laws. Eggs were housed in an incubator at 18°C until further use. They were then incubated at 38.5°C in a humidified incubator until the desired developmental stage. In all experiments, embryos were randomized in each experimental group and were harvested at embryonic day 4 (4 days post-fertilization).

### Cell lines

The A375P human melanoma cell line was obtained from ATCC and cultured in DMEM supplemented with 10% FBS (Cambrex) and 100 U/ml penicillin-streptomycin (Invitrogen). Stable expression of GFP in A375P was obtained by transduction of HIV1-based lentiviral particles as explained below.

Patient-derived short-term cultures (< 10) were established from a *BRAF^V600^* metastatic melanoma, before treatment for GLO, or after acquisition of resistance to vemurafenib for GLO-R. These short-term cell cultures were grown in RPMI complemented with 10% FBS and 100 U/ml penicillin-streptomycin.

### Viral infection and plasmid

Self-inactivating HIV1-derived vectors were produced by the lentivectors production facility / SFR BioSciences Gerland - Lyon Sud (UMS3444/US8) and encode the green fluorescent protein (GFP) under the control of a SFFV promoter (SIN-HIV-SFFV-eGFP). Briefly A375P cells were plated in six well plates (5×10^5^ cells per well) in complete medium. After 2 hours medium was replaced with 2 mL medium containing 2 % FBS and 2 mg/mL polybrene (Sigma). After an hour this medium was removed and replaced with 2mL of medium containing 5x 10^6^ IU of lentiviral vector. After 16 hours medium was removed and cells rinsed and incubated with normal medium (10% FCS). Analysis by FACS showed that close to 100% of cells were positive for GFP. Medium from semi-confluent transduced cells showed no capacity to transfer GFP expression to naive control cell lines, indicating that infectious viruses were not produced by the transduced cells.

### Human samples

Patient samples OF-MEL-001, OF-MEL-002, OF-MEL-003, OF-MEL-004, OF-MEL-005, OF-MEL-025, OF-MEL-026, OF-MEL-027, OF-MEL-028 and associated histological and clinical data were obtained from the Biological Resource Center of the Lyon Sud Hospital (Hospices Civils de Lyon) following surgery as their standard of care with patient’s informed signed consent to reuse biological samples for research purposes. Human melanoma sample OF-MEL-020 was obtained from NeuroBioTec (CRB HCL, Lyon France, Biobank BB-0033-00046) and is part of a collection registered at the French Department of Research (DC 2008-72).

Cutaneous melanoma biopsies OF-MEL-033, OF-MEL-034, OF-MEL-035 were obtained from patients included in the clinical trial NCT0439672, performed in the Lyon Sud Hospital (Hospices Civils de Lyon). Cutaneous biopsies were taken either from primary lesions or cutaneous metastases, and biopsied before treatment. Human tumor samples and clinical data were collected once the patients signed their informed consent to be included in the study. This minimally invasive study was approved by the national health authorities and ethics committee “Comité de Protection des Personnes Sud Méditerranée III” (n° ANSM 2019-A00900-57). Following surgery, resected tumors and biopsies were collected and stored in AqIX-RSI sterile medium (AqIX) for a maximum of 24 hours. All samples were cryopreserved prior to engraftment, except for OF-MEL-001 and OF-MEL-026 samples which were directly implanted in avian embryos as fresh samples.

### Human frozen samples

Tumors were washed with Ca^2+^,Mg^2+^-free phosphate-buffered saline (PBS) (Life Technologies), crushed with a sterile scalpel into small tissue pieces of 1 mm^3^, and put in freezing medium containing DMEM Glutamax medium (Gibco) supplemented with 10% FBS (Pan-biotech) and 10% DMSO (Sigma-Aldrich) in a cryotube prior to cryopreservation.

### *In ovo* xenografts of melanoma samples

Embryonated eggs were incubated at 38.5°C in a humidified incubator until HH14 stage. Fresh or frozen patient samples were dissociated in Hank’s Balanced Salt Solution (HBSS) with 156 units/mL of type IV collagenase, 200 mM CaCl2 and 50 units/mL DNase I for 20 minutes at 37°C and then incubated with 5 mg/mL trypsin for 2 minutes at 37°C under gentle mixing. Non-dissociated tissue was removed by filtration trough 0.4 μm nylon cell strainer (BD falcon). Non-fluorescent cell lines or patient samples were labeled with an 8 μM CFSE solution (Life Technologies). Stage HH12 chick embryos were grafted with fluorescent cells at the top of the dorsal neural tube within the migration staging area, with a glass capillary connected to a pneumatic PicoPump (PV820, World Precision Instruments) under a fluorescence stereomicroscope. For cell lines, approximately 2,500 living cells were grafted in each embryo, 200 to 300 for patient samples. For patient samples, the full cellular content obtained after dissociation was engrafted possibly including stromal and/or immune cells.

### Determination of drug maximum tolerated dose in chick embryos

Drugs were injected intravenously. Twenty-four hours after injection, chick embryos were harvested, weighed (Sartorius Quintix35-1S) and measured along the rostro-caudal axis using the Leica LASX image analysis software. The Body Surface Area (BSA) was calculated using Dubois & Dubois formula: BSA (m^2^) = 0.20247 × height (m)^0.725^ × weight (kg)^0.425^. The morphology / anatomy of each embryo was systematically analyzed to check their correct stage-related development. The criteria observed were: the survival (heart beating), the craniofacial morphology (presence of each cerebral compartment and eyes), the presence of four limb buds, the cardiac morphology, and the anatomy of embryonic annexes such as the allantois.

### Immunofluorescence on cryosections

Chick embryos were harvested and fixed in 4% Paraformaldehyde (PFA). Embryos were embedded in 7.5% gelatin - 15% sucrose in PBS to perform 20 μm transverse cryosections. Heat-induced epitope retrieval was performed by immersion in antigen unmasking solution (citrate buffer) at 70°C for 2 hours. Permeabilization and saturation of sections were performed in PBS - 3% Bovine Serum Albumin (BSA) - 0.5%. Triton. Anti-Ki67 (1/200, ab15580, Abcam), anti-MITF (1/100, clone C5, MAB3747, Merck-millipore), anti-HNK1 (1/50, clone 3H5, DSHB), anti-SOX10 (1/200, 89356, Cell Signaling) were applied to cryosections and incubated overnight at 4°C. Alexa 555 anti-rabbit IgG (1/500, A21429, Life Technologies), Alexa 555 anti-mouse IgG (1/500, A31570, Life Technologies), FluoProbes 647H donkey anti-mouse IgG (1/500, FPSC4110, Interchim) were used as secondary antibody. Nuclei were stained with Hoechst (H21486, Invitrogen). Slices were imaged with a confocal microscope (Olympus, FV1000, X81) using either a 10X objective for whole slice imaging or a 40X objective to focus on Ki67, MITF and SOX10 immunolabeling.

### Immunofluorescence analyses on paraffin-embedded samples

3-μm tissue sections were cut from PFA-fixed paraffin-embedded embryos. The sections underwent immunofluorescence staining using the OPAL™ technology (Akoya Biosciences) on a Leica Bond RX. A 7-color panel was designed. Anti-SOX10 (1/1000, Santa Cruz sc-365692) and ant-MITF (1/200, Sigma, 284M-96) primary antibodies were used. DAPI was used for nuclei detection. Sections were digitized with a Vectra Polaris scanner (Perkin Elmer, USA).

### Tissue clearing, whole mount SPIM imaging and image analysis

PFA-fixed HH25 embryos were cleared using an adapted Ethyl-Cinnamate protocol (Klingberg *et al*, 2017). Briefly, tissues were dehydrated in successive ethanol baths finally cleared in Ethyl Cinnamate (Sigma, 112372). Cleared samples were imaged using the UltraMicroscope SPIM (LaVision Biotech). 3D-images were built using Imaris™ software. Volumetric analysis was performed using the Imaris^™^ “Surface” module adjusted on CFSE or GFP fluorescence. The compaction index was determined as the ratio between the total volume occupied by tumor cells in an engrafted embryo and the number of fluorescent (CFSE^+^) objects segmented with the Imaris^™^ Surface module.

### *In vitro* cell survival assays

The CellTiter-Glo Luminescent Cell Viability Assay (ATP assay) (Promega) was used. 1,000 cells in 96-well plates were treated with three by 3-fold dilutions of the indicated drugs (BRAFi PLX4032, MEKi GDC-0973) for 72 hours in a final volume of 100 μL. Luminescence was measured (Tekan). Control wells with DMSO were used for normalization.

### Immunoblot analyses

Cells were washed twice with PBS containing CaCl_2_ and then lysed in a 100 mM NaCl, 1% NP40, 0.1% SDS, 50 mM Tris pH 8.0 RIPA buffer supplemented with a complete protease inhibitor cocktail (Roche, Mannheim, Germany) and phosphatase inhibitors (Sigma-Aldrich). Protein expression was examined by Western blot using the anti-ZEB1 (H102, 1/200, Santa Cruz), anti-ZEB2 (1/500, Sigma), anti-MITF (clone C5, ab80651, 1/500, Abcam), antibodies for primary detection. Loading was controlled using the anti-GAPDH (1/20,000, Millipore) antibody. Horseradish peroxidase-conjugated rabbit anti-mouse, goat anti-rabbit, and donkey anti-goat polyclonal antibodies (Dako, Glostrup, Denmark) were used as secondary antibodies. Western blot detections were conducted using the Luminol reagent (Santa Cruz).

### RNA-Seq

mARN from GLO and GLO-R were extracted in duplicates with the RNeasy mini kit (Qiagen) with DNase treatment. RNA libraries were prepared with the NextFlex Rapid Directional mRNA-Seq kit (Bioo-Scientific) with polyA+ mRNA enrichment, and sequenced on the ProfileXpert platform, on an Illumina Nextseq500 sequencing machine with a single read protocol (75bp; 30M reads). After demultiplexing and trimming, trimmed reads were mapped using TopHat 2.1.00b (Trapnell *et al*, 2009) against Human genome (hg19, GRCh37 Feb. 2009 from UCSC) in order to identify expressed genes. Reads mapping on each transcript were numbered and normalized using Cufflinks v.2.1.1 (Trapnell *et al*, 2010), fold change between the different groups were calculated using median of groups, and p-values were calculated using a t-test with equal variance and no p-value correction. Those calculations were performed using a proprietary R script. Mapped reads for each sample were counted and normalized using FPKM method (Fragments Per Kilobase of exon per Milion of mapped reads). Differentially expressed transcripts (|lFC| >1.5; p < 0.05) were analyzed between GLO versus GLO-R. Single sample GSEA (ssGSEA) scores were computed on FPKM normalized data through gsva R package.

## Data availability

The data reported in this paper are deposited in the Gene Expression Omnibus (GEO) database under the accession number GSE206689.

## Quantification and statistical analysis

Statistical treatment of data was performed with Prism 9.0e (GraphPad). For parametric tests, both normality and variance homoscedasticity were checked. All statistical tests were two-sided.

## The Paper Explained

### Problem

Metastatic melanoma patients carrying a BRAF^V600^ mutation can be treated with targeted therapies (BRAF and MEK inhibitors) but resistance occurs. Predicting patient response to targeted therapies is crucial to guide clinical decision, since these patients may also be directed to first-line immunotherapy. Mouse patient-derived xenograft (PDX) models are incompatible with personalized medicine approaches because of their long timeframe.

### Results

Herein, we developed a highly efficient patient-derived xenograft model using the avian embryo as a host (AVI-PDX^™^), enabling fast (few days) and reproducible tumor engraftment of melanoma patient samples, preserving key molecular and phenotypic features. We show that response to targeted therapies can be reliably modeled in these AVI-PDX^™^, making it a valuable preclinical tool for assessing efficacy of combination treatments in melanoma.

### Impact

We provide proof-of-concept that the AVI-PDX^™^ models the diversity of responses of melanoma patients to BRAFi/MEKi, within days, hence positioning it as a valuable tool for the design of personalized medicine assays.

## Acknowledgements

The authors would like to thank Séverine Croze and Joël Lachuer, from « ProfileXpert » (service de « Génomique & Microgénomique », Université Lyon 1, SFR santé LYON-EST, UCBL-INSERM US 7-CNRS UMS 3453) for RNA-Seq experiments. We also thank B. Manship for English editing.

YT was supported by a fellowship from Ligue Nationale contre le Cancer and MP by a fellowship from “Région Rhônes Alpes” and the Association pour la Recherche contre le Cancer (ARC). Oncofactory was supported by the program “Innovation clinique” of the Hospice Civils de Lyon-Lyonbiopôle.

## Contributions

L.J. performed the *in ovo* engrafting experiments, 3D imaging, image analyses and immunofluorescence studies on slices.

S.D. led the clinical aspects of the study (design of the study, patient sampling and processing).

C.C. participated to the *in ovo* engrafting experiments, 3D imaging and image analyses.

Y.T. did the *in vitro* experiments with GLO and GLO-R cells.

M.G. performed immunofluorescence staining of paraffin sections.

M.P. did the RNASeq experiments and analyses.

M.L. helped with the injection of therapies in avian embryos.

R.T. oversaw tumor volume analyses in avian embryos and management of the patient samples and associated data; corrected the manuscript.

M.LB. managed the writing, submission to ethics committees and follow up of the clinical assay.

D.M.-B. supervised the statistical analyses of data collected in the clinical assay.

A.E. supervised the collection of patient samples and associated clinical data; corrected the manuscript.

V.C. co-conceived the study and associated clinical assay, oversaw the work related to the setup of the AVI-PDX^™^ model.

J.C. co-conceived the study, oversaw aspects related to melanoma samples and cell lines, characterization of their phenotypes and wrote the manuscript.

C.D.-B. co-conceived the study and associated clinical assay, oversaw work and data related to the setup and use of the AVI-PDX^™^ model and wrote the manuscript.

All authors have contributed to the manuscript.

---

**Supplementary Table 1:** Characteristics of melanoma patient samples engrafted in avian embryos.

| Patient # | Sample characteristics |  |  |  |  |  | nb of grafted embryos | % tumor intake |
| --- | --- | --- | --- | --- | --- | --- | --- | --- |
|  | sex | nature | localization | stage | thickness | nb mitosis / mm<sup>2</sup> |  |  |
| Mel#1 | M | primitive | scalp | I | 0.40 mm | <1 | 11 | 18% |
| Mel#2 | M | primitive | foot arch | IV | 3.50 mm | 4 | 17 | 100% |
| Mel#3 | M | primitive | shoulder | II | 2.05 mm | 2 | 20 | 50% |
| Mel#4 | M | primitive | back | I | 0.50 mm | <1 | 12 | 83% |
| Mel#5 | M | primitive | back | IV | 5.00 mm | 13 | 23 | 83% |
| Mel#20 | M | metastasis | metastasis | IV | ND | ND | 76 | 100% |
| Mel#25 | M | primitive | back | IIC | 6.00 mm | 15 | 21 | 95% |
| Mel#26 | F | metastasis | behind ear | IIIB | ND | ND | 55 | 100% |
| Mel#27 | F | primitive | calf | II | 1.40 mm | <1 | 24 | 100% |
| Mel#28 | M | metastasis | knee | IV | ND | ND | 100 | 100% |
| Mel#33 | M | metastasis | leg | IV | ND | ND | 20 | 83% |
| Mel#34 | F | metastasis | arm | IV | ND | ND | 28 | 88% |
| Mel#35 | M | metastasis | back | IV | ND | ND | 25 | 100% |

**Supplementary Figure 1:**
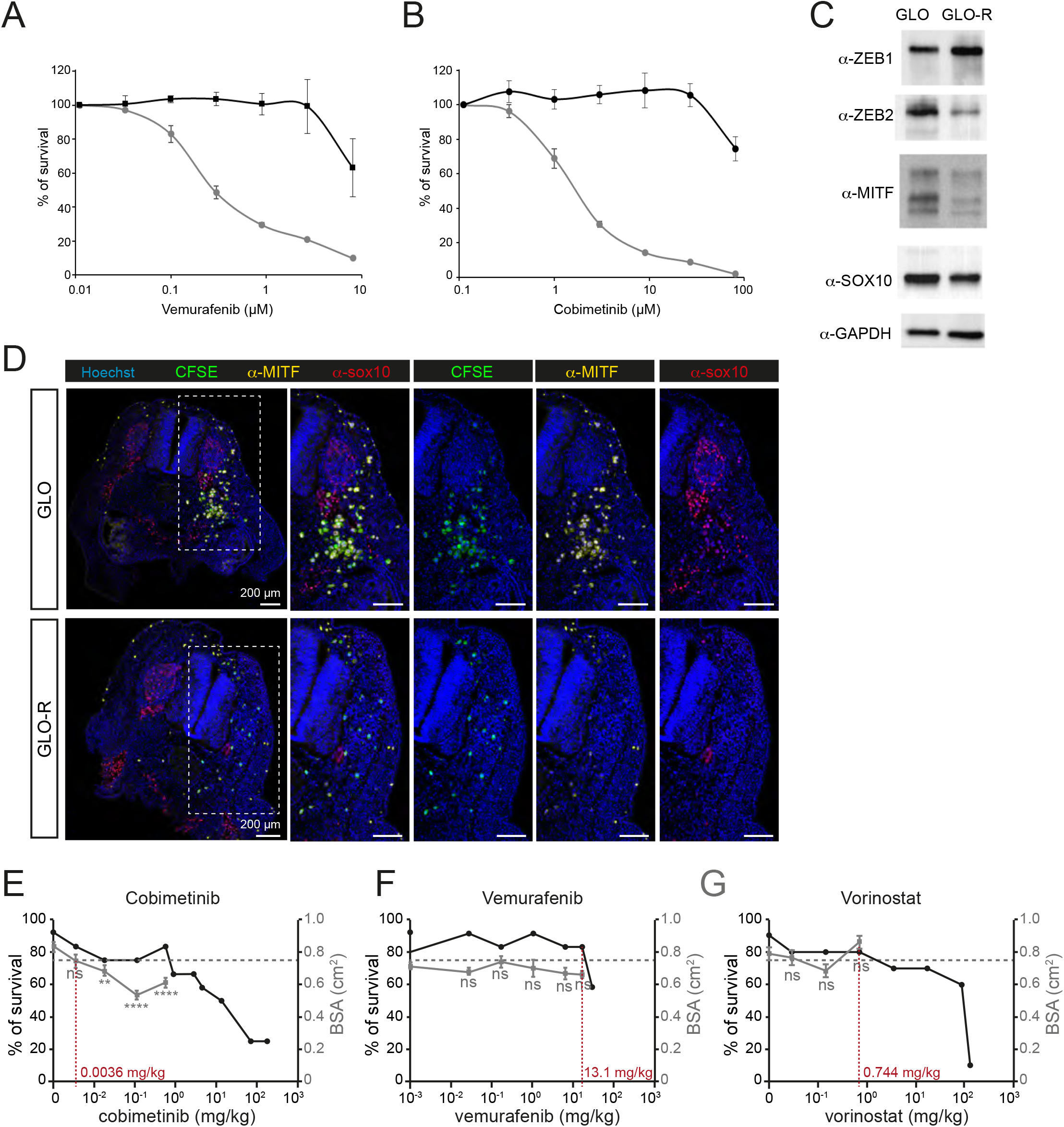
Characterization of GLO and GLO-R cell lines *in vitro* and in the AVI-PDX™ model. **A,B.** Survival rate of GLO and GLO-R cells upon exposure to increasing doses of Vemurafenib (A) or Cobimetinib (B) for 72 hours. **C.** Detection of ZEB1, ZEB2, MITF and SOX10 expression by Western blot in GLO and GLO-R cells, using GAPDH as a loading control. **D.** Immunofluorescent labelling of MITF (yellow) and SOX10 (red) in HH25 avian embryos engrafted with GLO or GLO-R cells, labelled with CFSE prior to the graft. Right panels are enlargements of the left panel for GLO and GLO-R grafts. **E-G**. Survival rate (left axis) and mean body surface area (BSA, right axis) of avian embryos injected with increasing doses of Vemurafenib (**E**), Cobimetinib (**F**) or Vorinostat (**G**). Each dose was administered to a minimum of 10 embryos, using excipient (NaCl) as a control. The maximum tolerated dose (MTD) was defined as the higher dose of drug associated with a survival rate higher than 75% and a mean BSA similar (*i.e*., non-statistically different) from embryos treated with NaCl. MTDs are indicated in red on the X-axis. Error bars indicate SEM. **: p < 0.01, ****: p < 0.0001, ns, non-significant using Student’s t-test compared with excipient.

